# Imprinted genes *Cdkn1c* and *Igf2* interact to promote terminal differentiation of adult NSCs

**DOI:** 10.1101/2022.01.24.477488

**Authors:** Anna Lozano-Ureña, Esteban Jiménez-Villalba, Laura Lázaro-Carot, Raquel Montalbán-Loro, Irene Martínez-Gurrea, Keiichi Nakayama, Martina Kirstein, Sacri R. Ferrón

## Abstract

Genomic imprinting is implicated in the control of gene dosage in neurogenic niches. Insulin-like growth factor 2 (*Igf2*), is an imprinted gene that shows biallelic expression only in the vascular compartment contributing to neural stem cells (NSCs) maintenance in the subventricular zone (SVZ) niche. The effects of this factor in adult NSCs differentiation are not well defined. We show here that IGF2 promotes cell fate commitment of NSCs by inducing the expression of another imprinted gene, the maternally expressed gene cyclin-dependent kinase inhibitor 1c (*Cdkn1c),* which encodes for p57 protein, eliciting cell cycle exit and terminal differentiation into astrocytes, neurons and oligodendrocytes. Using a conditional mouse model with *Cdkn1c*-deficient neural progenitors, we confirm that IGF2 and p57 interact in a common pathway to regulate the differentiation program of adult NSCs. This occurs through a mechanism involving the PI3K-Akt pathway that mediates regulation of p57 expression. We also show that the imprinted state of the *Cdkn1c* gene is not altered after IGF2 treatment confirming maternal expression of the gene in NSCs. Our results identify a molecular mechanism by which a paracrine factor produced and secreted by the neurogenic niche compartment can modulate *Cdkn1c* dosage to trigger differentiation of adult NSCs.

## Introduction

Adult neurogenesis in the subventricular zone (SVZ) is an orderly multistep process in which self-renewing astrocyte-like neural stem cells (NSCs or B1 cells) produce mature progeny via transit-amplifying progenitors (TAP or C cells) (Chaker et al., 2016). These progenitors rapidly divide to give rise to neuroblasts (A cells) that migrate to the olfactory bulb (OB) where they fully differentiate and integrate as interneurons (Calzolari et al., 2015; Chaker et al., 2016; Gotz et al., 2016; Ponti et al., 2013). The SVZ also gives rise to oligodendrocytes and astrocytes that integrate into the corpus callosum (CC) (Menn et al., 2006; Sohn et al., 2015). Sustained neurogenesis throughout adult life also occurs in the subgranular zone (SGZ) of the dentate gyrus (DG) in the hippocampus and is hypothesized to be involved in behavioural/cognitive processes such as memory and in diseases of the central nervous system (CNS) (Gage et al., 1998; Goncalves et al., 2016).

Genomic imprinting is an epigenetic process that causes genes to be expressed depending on their parental origin (John and Surani, 2000). A relatively small subset of genes within the mammalian genome (0.4%) is imprinted showing monoallelic expression in the whole organism or in specific tissues favoring the maternal or the paternal allele (John and Surani, 2000). Imprinted expression is initially determined by differential DNA methylation that is established in the germline (Surani, 1998). The prevalence of genomic imprinting is higher in the brain than in other organs and many genes with a crucial role in neurodevelopment are expressed in a maternal or paternal specific manner (Perez et al., 2016). For instance, *Igf2* is an imprinted gene that has been shown to regulate NSCs in the adult mouse SVZ (Ferron et al., 2015; Lehtinen et al., 2011; Ziegler et al., 2019). IGF2 is thought to control NSC pool by influencing its cell cycle, indeed IGF2 was shown to promote expansion of NSCs more potently than either insulin-growth-factor 1 (IGF1) or standard growth media (Alagappan et al., 2014; Ziegler et al., 2014; Ziegler et al., 2019). Detailed analysis of the SVZ niche has shown that IGF2 is secreted in a paracrine manner by the choroid plexus (CP) epithelium, where it is accessible to NSCs contacting the cerebrospinal fluid (CSF) (Bracko et al., 2012; Ferron et al., 2015; Lehtinen et al., 2011; Ziegler et al., 2014). The brain vasculature and leptomeninges also secrete IGF2 into the neurogenic niches (Ferron et al., 2015). Remarkably, *Igf2* is only expressed by the paternal allele in most tissues, but in these compartments the maternal allele is specifically activated resulting in biallelic expression of the gene (Ferron et al., 2015; Lehtinen et al., 2011).

*Cdkn1c* is another imprinted gene expressed only by the maternal allele (Pateras et al., 2009) and the misregulation of its expression has been associated with human growth disorders, such as Beckwith-Wiedemann and Silver-Russell syndromes, as well as with the onset of several types of cancers (Bastaki et al., 2016; Nakashima et al., 2015). *Cdkn1c* encodes for p57 protein and belongs to the cyclin-dependent kinase inhibitors CIP/KIP family, which also includes p21 and p27. p57 is the least studied CIP/KIP member although it has been demonstrated to play a fundamental role in regulating cell cycle and differentiation during mammalian development (Rossi et al., 2018). Indeed, studies in fetal stem cells such as cerebral cortical precursors showed that this cell cycle regulator controls multiple stages of corticogenesis such as cell cycle exit, differentiation and migration (Furutachi et al., 2015; Laukoter et al., 2020; Mairet-Coello et al., 2012; Tury et al., 2011). Importantly, p57 accumulates in MASH1+ proliferating telencephalic neural progenitors repressing neuronal differentiation independently of cell-cycle exit acting as a direct repressor of transcription (Joseph et al., 2009). p57 has also been shown to be a key factor during embryonic specification of slowly dividing NSCs as deletion of p57 impairs the emergence of adult NSCs in the SVZ (Furutachi et al., 2015). In the adult brain, an important function of p57 has also been demonstrated in NSCs derived from the SGZ of the mouse hippocampus, where its deletion abrogates quiescence and activates proliferation (Furutachi et al., 2013). Moreover, a detailed analysis of hematopoietic stem cells (HSCs) lacking p57 has confirmed the role of this cell cycle inhibitor as a key molecule in quiescence and self-renewal (Matsumoto et al., 2011; Tesio and Trumpp, 2011; Zou et al., 2011). This reveals the importance of its proper expression dosage during development, however, the potential role of p57 in the regulation of neurogenesis in the adult SVZ niche has not been addressed.

Studies in the hematopoietic system showed that IGF2 acts as a potent growth factor for adult bone marrow cells by stimulating their *ex vivo* expansion (Barroca et al., 2017; Zhang and Lodish, 2004) and a direct link between IGF2 and the cell cycle regulator p57 has been shown in this system (Thomas et al., 2016). In fact, overexpression of IGF2 in purified adult murine HSCs resulted in the upregulation of the *Cdkn1c* gene giving rise to an arrest of HSCs in the G0/G1 phase of the cell cycle and promoting the formation of multi-lineage colonies *in vitro* (Thomas et al., 2016). Nevertheless, the possible connection between IGF2 and p57 in the adult NSC population has not been studied.

In our study, we report that IGF2 promotes cell fate commitment by inducing the expression of *Cdkn1c* mRNA and p57 protein. This enhances terminal differentiation of adult NSCs *in vitro*. We also identify a methylation-independent mechanism of control of the *Cdkn1c* imprinted gene after IGF2 treatment, which involves the PI3K-Akt pathway during the differentiation of adult NSCs.

## Results

### IGF2 promotes terminal differentiation of neural stem cells

NSCs isolated from the adult SVZ were expanded in suspension as neurospheres in the presence of strong mitogenic stimulation with epidermal growth factor (EGF) and fibroblast growth factor 2 (FGF2) (Ferron et al., 2007). These proliferating neurospheres were disaggregated, plated on Matrigel (adherent assay), and cultured for 2 days *in vitro* (2 DIV) in medium containing only FGF2 to induce neural progenitors differentiation (Belenguer et al., 2016). Afterwards, all mitogens were withdrawn from the medium and it was supplemented with 2% fetal bovine serum (FBS) required for terminal differentiation (**Fig. 1a**). Under these conditions, differentiation is pushed forward and NSCs stop dividing and initiate an orderly program of commitment and differentiation into neurons, oligodendrocytes, and astrocytes during the following 5 days (7 DIV) (**Fig. 1a**).

**Figure 1.**
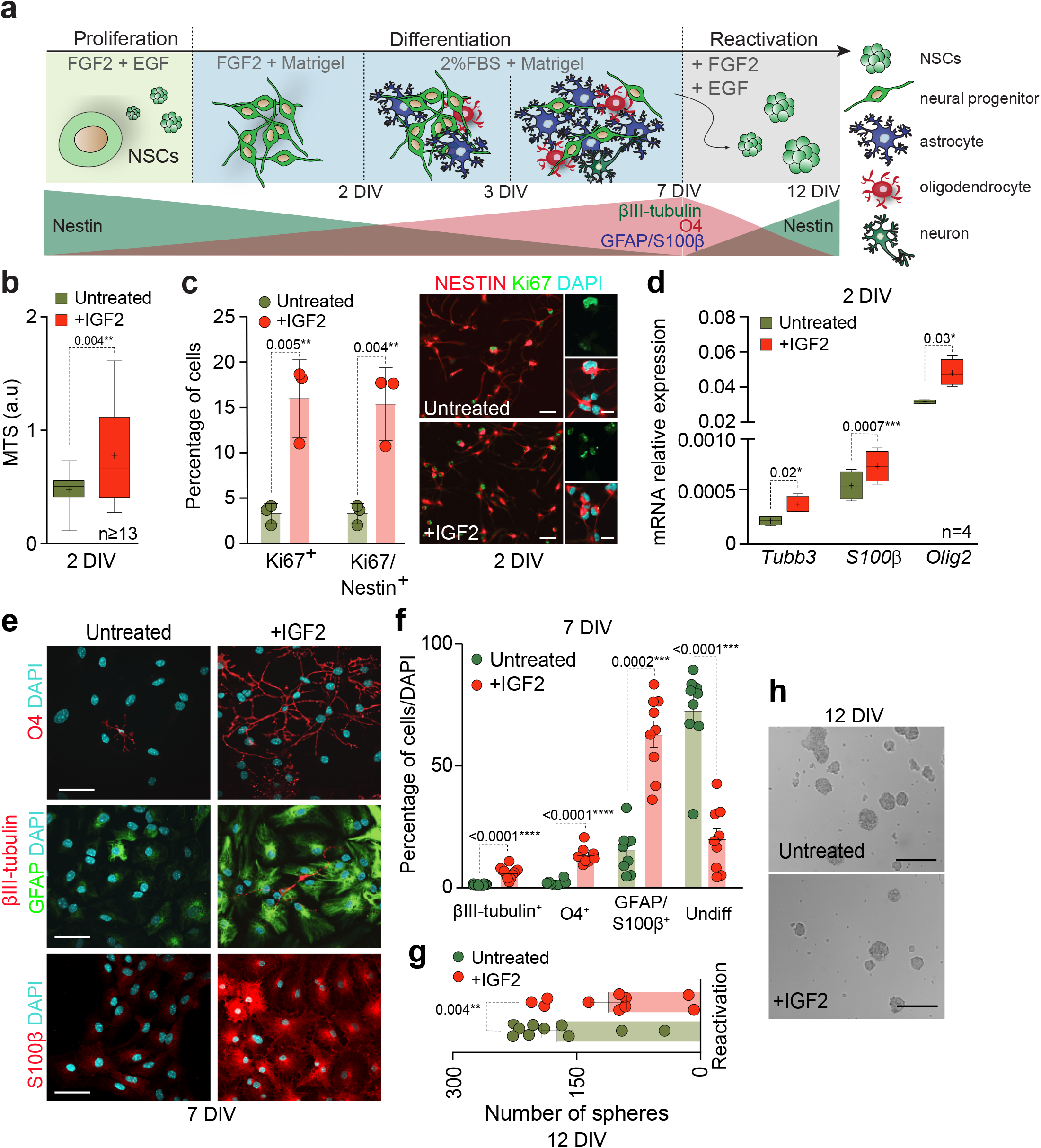
IGF2 promotes terminal differentiation of adult NSCs. **(a)** Schematic representation of differentiation and reactivation processes in adult NSCs. For expansion, cells were grown in presence of mitogens (EGF and FGF) and then cultured attached to Matrigel in absence of EGF for 2 days. For terminal differentiation, NSCs were cultured for 5 more days in presence of serum, and FGF was removed from the medium. For reactivation, NSCs were detached and cultured in proliferation promoting conditions. **(b)** MTS assay performed after 2 days (2 DIV) of differentiation in the presence or absence of IGF2 (100 ng ml^−1^). Measured as arbitrary units (a.u.). **(c)** Percentage of cells positive for Nestin and Ki67 after 2 days (2 DIV) of differentiation in the presence or absence of IGF2 (left panel). Immunocytochemistry images for Nestin (red) and Ki67 (green) in untreated and IGF2-treated cultures are also shown (right panel). **(d)** Quantitative PCR (qPCR) for *βIII-tubulin (Tubb3)*, *S100β* and *Olig2* genes in NSCs after 2 days in differentiation conditions. *Gapdh* was used as housekeeping gene. **(e)** Immunocytochemistry images for O4 (red), GFAP (green) and βIII-tubulin (red) in NSCs after 7 days of differentiation in presence or absence of IGF2 (upper and middle panels). Immunocytochemistry images for S100β in untreated and IGF2-treated cultures are also shown (lower panels). **(f)** Percentage of cells positive for βIII-tubulin, O4 and GFAP/S100β and in untreated and IGF2-treated cultures after 7 days in differentiation-promoting conditions. The percentage of undifferentiated cells is also determined. **(g)** Number of spheres formed after reactivation by detaching 7 days-differentiated NSCs and replating them for 5 more days in proliferation-promoting conditions (FGF and EGF) in presence or absence of IGF2 (left panel). **(h)** Representative images of neurospheres formed by differentiated cultures in the presence or absence of IGF2 (right panel). DAPI was used to counterstain DNA. All error bars show s.e.m. of at least 3 cultures per condition. P-values and number of samples are indicated. Scale bars in c and e: 30 μm; in h: 100 μm.

IGF2 plays an important role in the regulation of adult neurogenesis, affecting NSCs proliferation capacity (Bracko et al., 2012; Ferron et al., 2015; Lehtinen et al., 2011). In order to study the effects of this factor in neural differentiation, NSCs from the adult SVZ were expanded and differentiated into the three neural lineages of the central nervous system in the presence of IGF2 (100 ng ml^−1^) (Belenguer et al., 2016). After 2 DIV of differentiation conditions, IGF2 treated NSCs showed an increase in proliferation, as indicated the MTS assay (**Fig. 1b**). Consistently, immunodetection of the proliferation marker Ki67, revealed an increase in the proportion of Nestin+ progenitors that were still proliferating after 2 DIV of differentiation in IGF2 treated cultures (**Fig. 1c**). Moreover, although the levels of expression of the gene *Nestin* were not altered in the presence of IGF2 (**Fig. S1a**), the expression of the neuronal gene *βIII-tubulin (Tubb3)*, the astrocytic gene *S100β* and the oligodendrocytic gene *Olig2* was increased in 2 DIV treated cells (**Fig. 1d**). Accordingly, culturing NSCs in the presence of IGF2 promoted their terminal differentiation into the three neural lineages, increasing the percentage of neurons measured as βIII-tubulin^+^ cells, astrocytes measured as cells expressing high levels of GFAP and oligodendrocytes measured as O4^+^ cells formed after 7 DIV of differentiation (**Fig. 1e, f**). Moreover, higher proportions of cells that were strongly positive for S100β, a protein largely absent from neurogenic GFAP^+^ cells (Raponi et al., 2007), were also observed after 7 DIV of differentiation in the presence of IGF2 (**Fig. 1e, f**) suggesting that IGF2 directly promotes proliferation and terminal differentiation of multipotent NSCs. To confirm whether enhanced differentiation in the presence of IGF2 was accompanied by a reduction in the capacity of differentiated NSCs to form neurospheres, 7-DIV differentiated adult NSCs cultures were detached and replated again in proliferating conditions (**Fig. 1a**). This led to the reactivation of a small proportion of cells with the capacity to still form neurospheres in non-adherent conditions after 5 more days *in vitro* (12 DIV) (**Fig. 1g, h**). As expected, the higher presence of S100β was consistent with a reduction in the neurosphere formation capacity in IGF2-treated NSCs cultures (**Fig. 1g, h**), indicating that the bias toward a more differentiated phenotype in the presence of IGF2 correlated with a reduction in stemness.

### IGF2 induces the expression of the maternally expressed gene *Cdkn1c* in differentiating NSCs

A direct link between IGF2 and the cell cycle regulator p57 has been shown in the hematopoietic system (Thomas et al., 2016). In fact, overexpression of IGF2 in purified adult murine HSCs resulted in the upregulation of the *Cdkn1c* gene giving rise to an arrest of HSCs in the G_0_/G_1_ phase of the cell cycle and promoting the formation of multi-lineage colonies *in vitro*. Neurosphere cultures grown in suspension under proliferating conditions exhibited low levels of *Cdkn1c* mRNA, but mitogen withdrawal induced a 4-fold increase in *Cdkn1c* gene expression as measured 2, 3 or 7 DIV later (**Fig. 2a**). Moreover, a significant increase in the percentage of cells with detectable levels of the p57 protein in differentiating cultures inversely correlated with the proportion of proliferating Ki67^+^ cells (**Fig. S1b**), suggesting that increased expression of p57 in the absence of mitogens might correlate with cell cycle exit and differentiation of adult NSCs. Accordingly, p57 protein was detected in Nestin^+^ progenitors after 2 and 3 days of *in vitro* differentiation and in GFAP^+^ astrocytes at 7 DIV (**Fig. 2b**).

**Figure 2.**
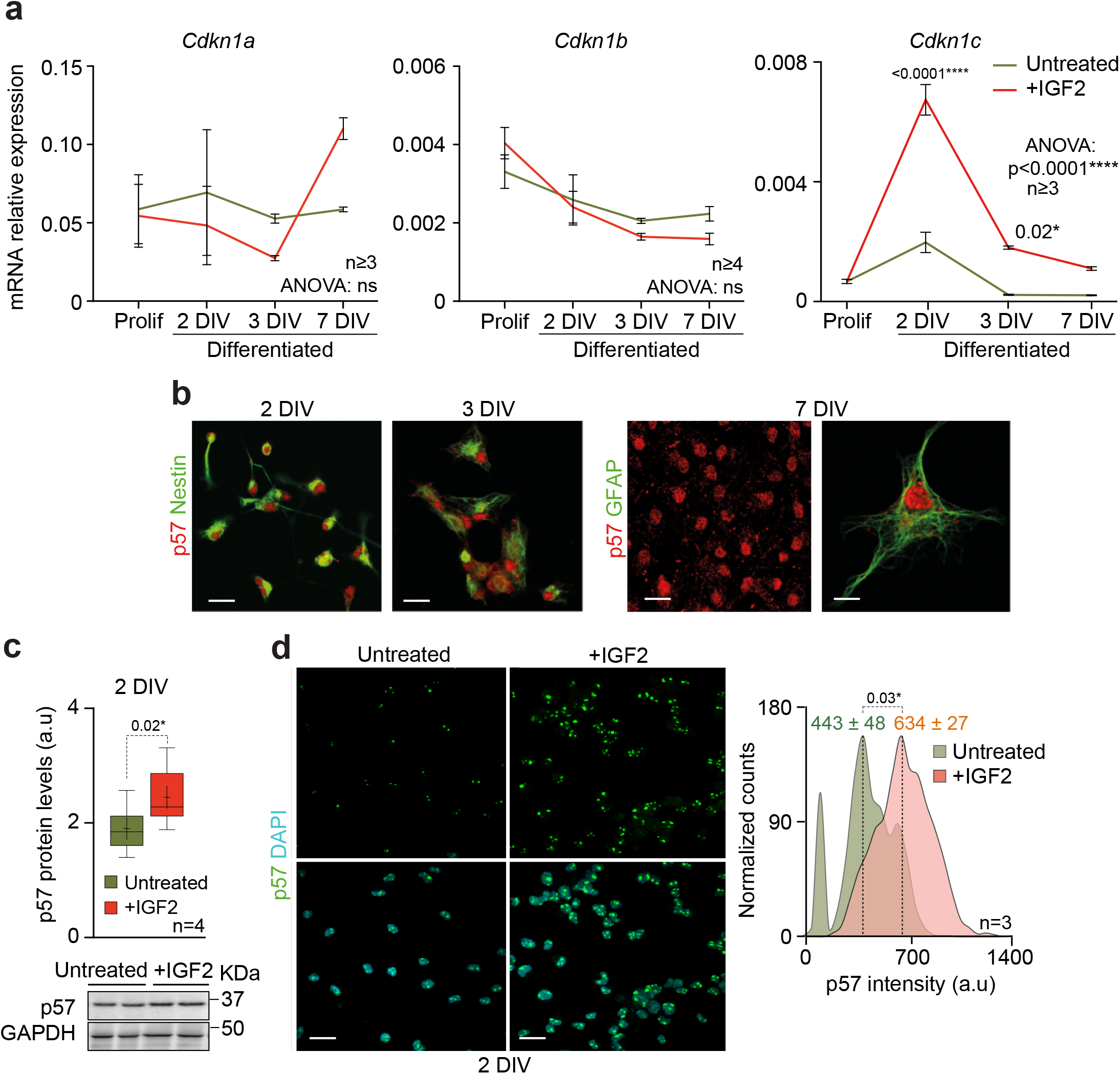
IGF2 induces *Cdkn1c* expression in adult NSCs in differentiation conditions. **(a)** qPCR for *Cdkn1a, Cdkn1b* and *Cdkn1c* in proliferating NSCs and after 2, 3 and 7 days in differentiation-promoting conditions in absence (green) or presence of IFG2 (red). *Gapdh* was used as a housekeeping gene. **(b)** Immunocytochemistry images for p57 (red), Nestin (green) or GFAP (green) in wild-type cultures in differentiation conditions. **(c)** Western-blot (WB) of p57 after 2 DIV of differentiation in untreated and IGF-treated cultures (lower panel). Quantification of p57 protein levels by WB (upper panel). **(d)** Immunochemistry for p57 (green) in untreated and IGF2-treated cultures after 2 days in differentiation-promoting conditions (left panel). Histograms showing p57 intensity (in arbitrary units, a.u) in wild-type cultures in the presence or absence of IGF2 after 2 DIV of differentiation. Mean intensities are indicated as dashed lines (right panel). DAPI was used to counterstain DNA. All error bars show s.e.m. of at least 3 cultures per condition. P-values and number of samples are indicated. Scale bars in b: 30 μm (high magnification image, 15 μm); in d: 50 μm.

To define the potential role of IGF2 in regulating *Cdkn1c* expression during adult NSCs differentiation a similar analysis of *Cdkn1c* mRNA levels in NSCs was performed after IGF2 treatment. This study revealed no changes in the levels of *Cdkn1c* expression, in proliferating conditions in the presence of IGF2 (**Fig. 2a**). However, *Cdkn1c* levels were significantly increased in differentiation-promoting conditions showing a maximum difference at the first step (2 DIV) of the differentiation process in the presence of IGF2 (**Fig. 2a**). No changes in the levels of expression of the other two members of the CKIs family of proteins, *Cdkn1a* and *Cdkn1b,* were observed at the same time of differentiation (**Fig. 2a**). Immunoblot detection for p57, confirmed the increased levels of the P57 protein in IGF2-treated cultures after 2 DIV of differentiation (**Fig. 2c**), consistent with the increased immunofluorescent detection of nuclear p57 in differentiation conditions after IGF2 treatment (**Fig. 2d**).

### IGF2 regulates *Cdkn1c* expression during NSCs differentiation through Akt activation

*Cdkn1c* expression is finely regulated by several epigenetic mechanisms including genomic imprinting. The mouse *Cdkn1c* gene belongs to the *Kcnq1ot1* imprinted cluster on mouse chromosome 7 and is canonically expressed from the maternally inherited chromosome (**Fig. S2a**) (Stampone et al., 2018). Therefore, we next explored whether the upregulation of *Cdkn1c* in IGF2-treated NSCs could be caused by a loss of imprinting of the *Cdkn1c* gene and thus acquisition of paternal expression of the gene. To test this, we assayed *Cdkn1c* imprinting state in NSCs at 2 DIV of differentiation in wild-type adult F1 hybrid offspring from reciprocal crosses of *Mus musculus domesticus (*C57BL6/J) and *Mus musculus castaneus* (CAST/EiJ) strains in which a single-nucleotide polymorphism (SNP) was identified at the *Cdkn1c* gene between the two subspecies (**Fig. S2b**). IGF2-treated and untreated cultures showed the expected maternally inherited imprinted expression of *Cdkn1c* (**Fig. 3a**), indicating that genomic imprinting was not altered after IGF2 treatment. The *Kcnq1ot1* imprinted cluster has a germline-derived differentially methylated region (gDMR) that is associated with imprinted expression of the genes within the cluster (**Fig. S2a**) (Hatada and Mukai, 1995; Stampone et al., 2018). Moreover, the *Cdkn1c* gene has a somatic DMR (sDMR) located at the promoter of the gene that also regulates its expression (Pateras et al., 2009; Stampone et al., 2018). Given that the imprinting state of the gene was not altered, and to further analyse whether DNA methylation of *Cdkn1c* promoter might be associated with the regulation of the expression of the gene after IGF2 treatment, we next determined the methylation levels of this sDMR by bisulfite sequencing. Consistently, IGF2-treated NSCs showed the expected levels of methylation (mean methylation percentage in Untreated: 61,6 ± 3,5 % and in +IGF2: 61,5 ± 3,6 %; p-value = 0.98) (**Fig. 3b**) supporting a methylation-independent function of IGF2 on the induction of *Cdkn1c* gene expression.

**Figure 3.**
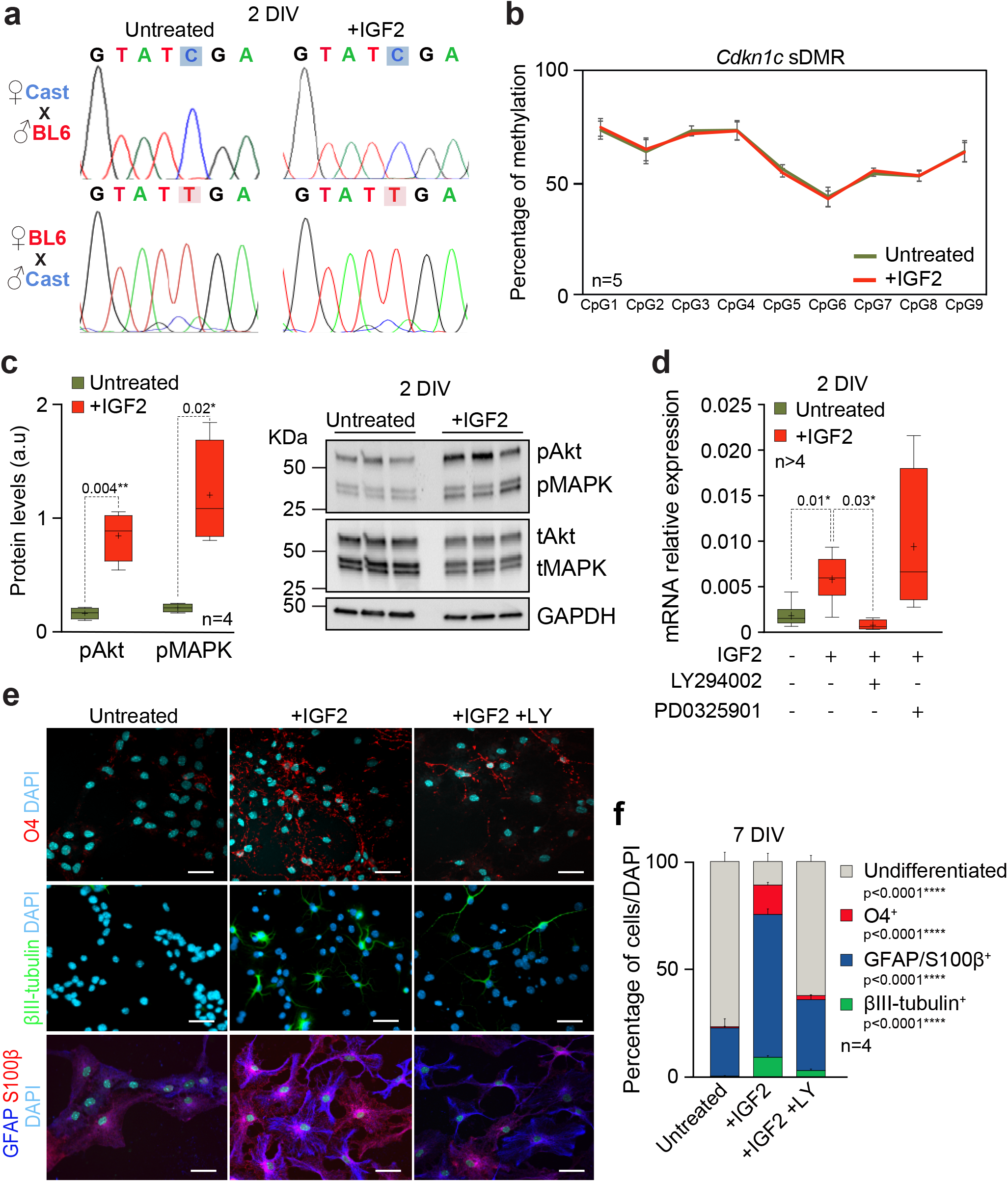
IGF2 promotes *Cdkn1c* expression in NSCs through a mechanism involving PI3K-Akt. **(a)** Sequence analysis of RT-PCR products from adult NSCs derived from reciprocal F1 hybrids mice from *Mus musculus domesticus* (abbreviated BL6) and *Mus musculus castaneus* (abbreviated Cast) after being treated or not with IGF2 in differentiation conditions. **(b)** Percentage of methylation determined by bisulfite sequencing and pyrosequencing at 9 CpG sites within the somatic DMR (sDMR) of *Cdkn1c* gene in untreated (green) or IGF2-treated (red) after 2 DIV of differentiation. All error bars show s.e.m. of at least 3 cultures per condition. **(c)** Quantification of phosphorylated forms of AKT and MAPK protein levels by WB (left panel). Data are presented relative to total Akt or MAPK protein. WB for phosphorylated AKT (pAkt) and MAPK (pMAPK) and total AKT (tAkt) and MAPK (tMAPK) in untreated and IGF2-treated cultures after 2 days in differentiation-promoting conditions (right panel). **(d)** qPCR for *Cdkn1c* in wild-type cultures in the absence or presence of IGF2 that have been additionally treated with the PI3K inhibitor LY294002 or the ERK1/2 inhibitor PD0325901 after 2 DIV of differentiation. **(e)** Immunocytochemistry images for O4 (red), βIII-tubulin (green), GFAP (blue) and S100β (red) in *wild-type* NSCs after 7 DIV of differentiation in the presence or absence of IGF2 that have been additionally treated with the PI3K inhibitor LY294002. **(f)** Percentage of cells that were positive for O4, GFAP/S100β and βIII-tubulin after 7 days in differentiation-promoting conditions in the presence or absence of IGF2 and that were additionally treated with LY. DAPI was used to counterstain DNA. All error bars show s.e.m. of at least 3 cultures per condition. P-values and number of samples are indicated. Scale bars in e: 30 μm.

It has been previously shown that insulin-like growth factors activate phosphatidylinositol 3-kinase/protein kinase B (also known as PI3K-Akt) and mitogen-activated protein kinase (MAPK) to promote cell cycle progression and differentiation (Bracko et al., 2012; Chirivella et al., 2017; Mairet-Coello et al., 2012). Precisely, the link between IGF2 and p57 has been shown to be dependent on the activation of PI3K/Akt pathway in HSCs (Thomas et al., 2016). Therefore, to elucidate the intracellular pathways downstream of IGF2 in differentiating NSCs, levels of phosphorylated Akt (pAkt) and ERK1/2 (pMAPK) expression in adult NSCs were determined after IGF2 treatment in differentiation conditions. This study revealed a significant increase of both pAkt and pMAPK in IGF2-treated NSCs after 2 DIV of *in vitro* differentiation (**Fig. 3c**). Interestingly, simultaneous treatment of adult NSCs cultures with IGF2 and the PI3K pharmacological inhibitor LY294002 blocked the effects of IGF2 on *Cdkn1c* expression (**Fig. 3d** and **Table S1**), whereas treatment with the ERK1/2 inhibitor PD0325901 did not modify the increased levels of *Cdkn1c* expression induced by IGF2 (**Fig. 3d**). These results demonstrated that IGF2 promoted *Cdkn1c* expression in NSCs through a mechanism involving PI3K-Akt but not ERK1/2-MAPK pathway.

To determine whether differentiation effects of IGF2 via p57 are mediated by activation of the PI3K-Akt pathway, wild-type NSCs were differentiated for 7 DIV into astrocytes, oligodendrocytes and neurons in the presence of IGF2 and the PI3K inhibitor LY294002. As previously described, IGF2 promoted an increase of the percentage of βIII-tubulin^+^ neurons, GFAP^+^/S100β^+^ astrocytes and O4^+^ oligodendrocytes cells (**Fig. 3e, f** and **Table S1**). However, the addition of LY294002 in the IGF2 treated cultures reverted the differentiation effects of the factor, resulting in cultures with less proportion of neurons, astrocytes and oligodendrocytes (**Fig. 3e, f** and **Table S1**). Accordingly, the proportion of undifferentiated cells was increased in the presence of LY294002 (**Fig. 3e, f** and **Table S1**). These data confirm that the PI3K-Akt pathway mediates regulation of *Cdkn1c* expression by IGF2 during the differentiation process of adult NSCs *in vitro*.

### p57 mediates the differentiation effects of IGF2 in adult NSCs

To finally evaluate whether p57 could mediate the differentiation effects of IGF2 in adult SVZ NSCs, a murine genetic model was generated by crossing mice carrying *loxP* sites flanking exons 2 to 4 of *Cdkn1c* gene (*Cdkn1c*^*loxp/loxp*^) (Matsumoto et al., 2011) with mice expressing the Cre-recombinase under the control of the Gfap promoter (*Gfap-cre*^*+/0*^) (**Fig. S3a**) (Garcia et al., 2004; Montalban-Loro et al., 2019). Given that, *Cdkn1c* is an imprinted gene expressed only from the maternal allele; these crosses were done reciprocally to generate heterozygous mice with either the maternal (*Cdkn1c-Gfap*^*mat*^) or paternal (*Cdkn1c-Gfap*^*pat*^) deficient allele (**Fig. S3a**). A significant reduction of *Cdkn1c* mRNA and protein was observed in 2 DIV *Cdkn1c-Gfap*^*mat*^ differentiating cultures in presence of IGF2 (**Fig. 4a, b**), however, no downregulation of the *Cdkn1c* gene was detectable in *Cdkn1c-Gfap*^*pat*^ heterozygous cultures (**Fig. 4a**) coincident with the canonical monoallelic expression of the gene. Importantly, the other two members of the CKIs family of proteins *Cdkn1a* and *Cdkn1b* were unaltered in *Cdkn1c-Gfap*^*mat*^ and *Cdkn1c-Gfap*^*pat*^ cultures (**Fig. S3b**).

**Figure 4.**
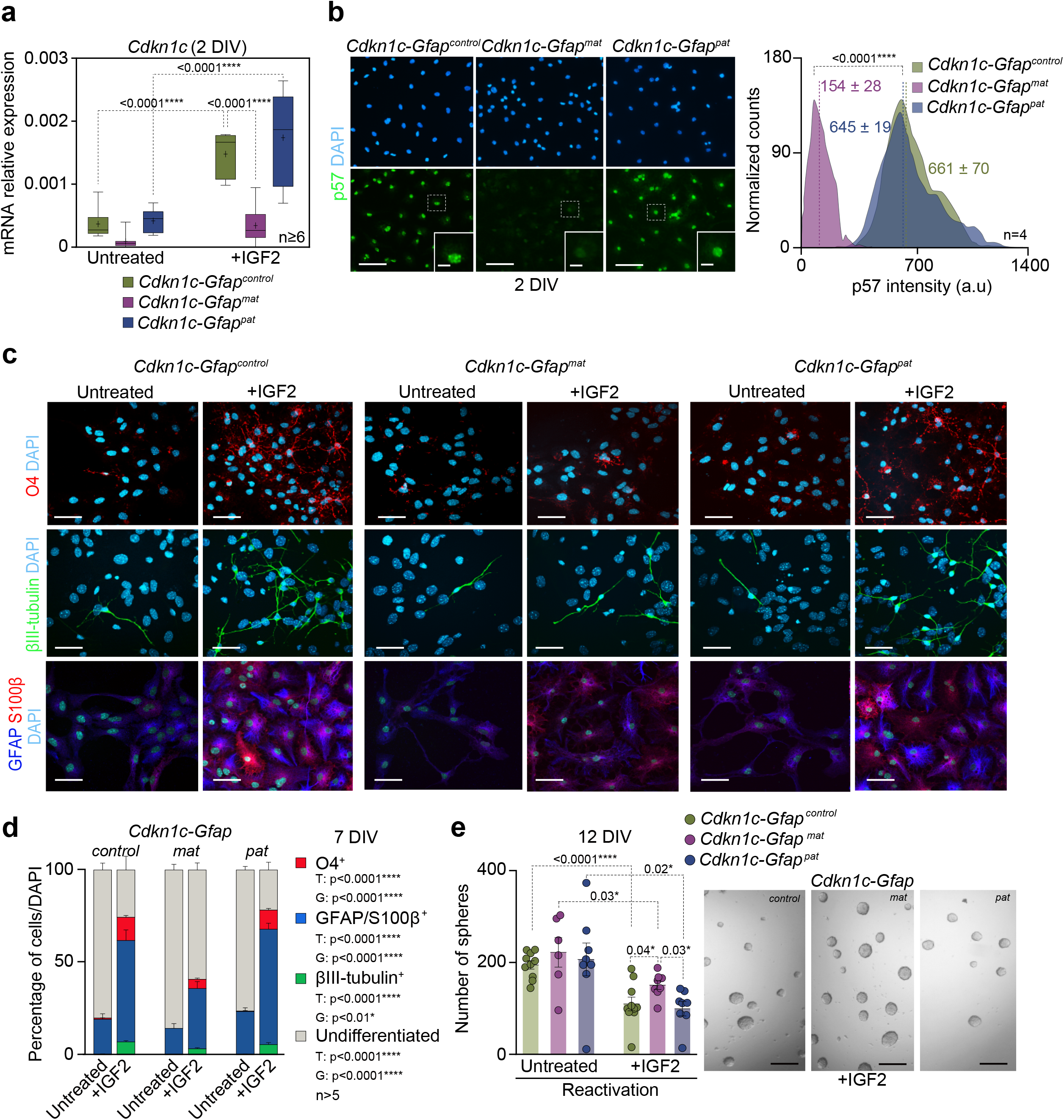
P57 mediates the effects of IGF2 on NSCs differentiation. **(a)** qPCR for the *Cdkn1c* gene in *Cdkn1c-Gfap*^*control*^, *Cdkn1c-Gfap*^*mat*^ and *Cdkn1c-Gfap*^*pat*^ NSCs after 2 DIV of differentiation in presence or absence of IGF2. *Gapdh* was used as a housekeeping gene. **(b)** Immunocytochemistry images for p57 (green) in *Cdkn1c-Gfap*^*control*^, *Cdkn1c-Gfap*^*mat*^ and *Cdkn1c-Gfap*^*pat*^ NSCs after 2 DIV of differentiation (left panel). Histogram showing p57 intensity (in arbitrary units, a.u) in *Cdkn1c-Gfap*^*control*^, *Cdkn1c-Gfap*^*mat*^ and *Cdkn1c-Gfap*^*pat*^ cultures after 2 DIV of differentiation. Mean intensities are indicated as dashed lines (right panel). **(c)** Immunochemistry images for O4 (red), βIII-tubulin (green), GFAP (blue) and S100β (red) in *Cdkn1c-Gfap*^*control*^, *Cdkn1c-Gfap*^*mat*^ and *Cdkn1c-Gfap*^*pat*^ NSCs after 7 DIV of differentiation and in the presence or absence of IGF2. **(d)** Percentage of cells that were positive for O4, GFAP/S100β and βIII-tubulin in untreated and IGF2-treated (100 ng ml^−1^) cultures after 7 days in differentiation-promoting conditions. **(e)** Number of neurospheres formed from *Cdkn1c-Gfap*^*control*^, *Cdkn1c-Gfap*^*mat*^ and *Cdkn1c-Gfap*^*pat*^ after detaching 7 DIV-differentiated NSCs in presence or absence of IGF2 and replating in proliferation conditions (left panel). Representative images of neurospheres formed by differentiated cultures in the presence or absence of IGF2 (right panel). DAPI was used to counterstain DNA. All error bars show s.e.m. of at least 4 cultures per condition. P-values and number of samples are indicated. Scale bars in b, c: 30 μm (inserts in b: 10 μm); in e: 100 μm.

Consistent with previously shown data, treatment of *Cdkn1c-*deficient cultures with recombinant IGF2 revealed that the *Cdkn1c* gene was significantly increased in *Cdkn1c-Gfap*^*pat*^ and *Cdkn1c-Gfap*^*control*^ after differentiation induction but its expression remained unchanged in *Cdkn1c-Gfap*^*mat*^ cultures (**Fig. 4a**). Therefore, to investigate whether the differentiation effects of IGF2 on adult NSCs might be mediated by p57, NSCs from the adult SVZ of heterozygous *Cdkn1c-Gfap*^*mat*^ and *Cdkn1c-Gfap*^*pat*^ and control mice were expanded and differentiated for 7 DIV into astrocytes, oligodendrocytes and neurons in the presence of IGF2. Again we found that IGF2 promoted terminal differentiation of NSCs increasing the percentage of βIII-tubulin^+^ neurons, GFAP^+^/S100β^+^ astrocytes and O4^+^ oligodendrocytes in control and *Cdkn1c-Gfap*^*pat*^ cultures (**Fig. 4c, d** and **Table S2**). However, the proportions of these three neural lineages in *Cdkn1c-Gfap*^*mat*^ cultures were not equally modified than wild-type in the presence of IGF2 (**Fig. 4c, d** and **Table S2**). Accordingly, the proportion of cells with the capacity to still form neurospheres in non-adherent cultures after differentiation was again reduced after IGF2 treatment in wild-type and *Cdkn1c-Gfap*^*pat*^ but not in *Cdkn1c-Gfap*^*mat*^ cultures (**Fig. 4e**). All these data confirm that maternal expression of *Cdkn1c* mediated the differentiation effects of IGF2 in adult NSCs *in vitro*.

## Discussion

This study shows that the imprinted genes *Igf2* and *Cdkn1c* interact to regulate the differentiation potential of adult NSCs *in vitro*. Treatment of NSCs with recombinant IGF2 promotes a more differentiated phenotype of NSCs and a reduction of their stemness *in vitro*. The promotion of terminal differentiation of NSCs correlates, with the induction of an increased level of the maternally expressed gene *Cdkn1c.* Consistently, differentiation of *Cdkn1c*-deficient neural progenitors in the presence of IGF2 shows a defective induction of cell fate, demonstrating that p57 mediates partially the differentiation effects of this niche factor. Moreover, although the imprinting state of *Cdkn1c* is not altered, IGF2 and p57 seem to interplay in a common pathway involving Akt-PI3K to control the differentiation programs of adult NSCs into astrocytes, oligodendrocytes and neurons (**Fig. S4**).

The formation of new neurons in the adult brain during adulthood is based on the balance between self-renewal and differentiation of neural progenitors into mature progeny (Calzolari et al., 2015; Chaker et al., 2016). The SVZ niche is complex and elegantly organized to favor access to the cerebral microvasculature and to CSF stimulating the growth and function of NSCs in an IGF2-dependent manner binding to type B1 cells primary cilium protruding from the wall of the lateral ventricle (Mirzadeh et al., 2008; Tavazoie et al., 2008). In the majority of tissues, particularly during development, *Igf2* expression occurs solely from the paternal allele. However we have previously described that *Igf2* is not only expressed from the paternal but also from the maternally inherited allele in the vasculature, CP, meninges and leptomeninges of the adult brain, and that such biallelic paracrine expression is required to support SVZ neurogenesis (Ferron et al., 2015). In contrast, NSCs within the SGZ rely on canonically imprinted autocrine IGF2 expressed only from the paternal allele to regulate hippocampal stem cell function (Ferron et al., 2015).

Although the effects of IGF2 in proliferation and survival of neural stem cells have been reported extensively for both late fetal and postnatal progenitors (Alagappan et al., 2014; Ferron et al., 2015; Lehtinen et al., 2011; Ziegler et al., 2014; Ziegler et al., 2019), the mechanisms by which IGF2 governs neurogenesis remain incompletely characterized. P57 has also been described to induce cell cycle exit, differentiation, neurite outgrowth and migration during embryonic cortical development (Imaizumi et al., 2020; Laukoter et al., 2020; Mairet-Coello et al., 2012) and is known to control NSCs quiescence in hippocampal neurogenesis (Furutachi et al., 2013). In the hematopoietic system, it has been previously shown that IGF2 upregulates expression of p57, through activation of the PI3K-Akt pathway. This increase of p57 allows the maintenance of the HSCs (Thomas et al., 2016). Indeed, loss of p57 in HSCs results in a decrease retention in G_0_ and impaired self-renewal potential (Thomas et al., 2016), illustrating the requirement for p57 to ensure maintenance of a quiescent, self-renewing population of HSCs. In the SVZ, IGF2 treatment also induces the expression of *Cdkn1c* and thus p57 protein. However, this IGF2-mediated effect occurs after mitogen withdrawal and not in proliferation conditions, promoting the terminal differentiation of NSCs.

Genomic imprinting controls the allelic expression of a subset of dosage-sensitive genes in a parent-of origin-dependent manner. Imprinted genes can be classified into canonical imprinted genes and genes that switch detectable expression of both parental alleles in different tissues and/or during development (Laukoter et al., 2020). Genes belonging to the second category include *Igf2*, which it has been shown to be biallelicaly expressed in the CP and vasculature in the SVZ niche having a critical function for postnatal neurogenesis (Ferron et al., 2015). Along the same lines, we have previously shown that loss of genomic imprinting of the paternally expressed gene *Dlk1*, is necessary to maintain NSC function in both neurogenic niches the SVZ and the SGZ (Ferron et al., 2011; Montalban-Loro et al., 2021). Thus, dynamic changes in genomic imprinting translate into biologically relevant functions. Notably, many of these imprinted genes work co-ordinately and even antagonistically to control fetal growth (Daniel et al., 2015; Hoffmann et al., 2014). Our expression data here show that the increased levels of expression of the imprinted gene *Cdkn1c* by the presence of IGF2 in the culture media occurs independently and without altering the imprinting state of the gene. Indeed, our study present an imprinting-independent model of regulation of *Cdkn1c* in which IGF2 upregulates expression of *Cdkn1c* through activation of the PI3K-Akt pathway. This IGF2-mediated increase in the levels of p57 *in vitro* promotes neuronal and glial differentiation of NSCs illustrating the requirement of the correct interaction of the two imprinted genes *Igf2* and *Cdkn1c* to ensure terminal differentiation of NSCs from the adult SVZ neurogenic niche.

Disturbed imprinted gene expression is associated with human disease (Monk et al., 2019; Tucci et al., 2019). For example, the human imprinted gene *CDKN1C* is the most frequently silenced or mutated gene in the imprinting genetic disorder Beckwith-Wiedemann syndrome (BWS) and aberrant imprinting of *IGF2* has also been shown to interact with CDK1C in this imprinting disorder (Caspary et al., 1999; Grandjean et al., 2000). Interestingly, both genes locate in the same imprinted gene cluster on human chromosome II (Grandjean et al., 2000). Moreover, alterations in both imprinted genes can lead to malignant cell transformation in different cancers (Bastaki et al., 2016; Nakashima et al., 2015). Therefore, our study highlights the importance to probe the relative contributions of imprinted genes to the differentiation potential of adult NSCs during the neurogenesis process and in non-pathological conditions. In future studies it will be important to investigate the precise roles of these genes in neurodevelopmental disease conditions.

## Methods

### Animals and *in vivo* manipulations

All transgenic mice used in the study were maintained in the C57BL6 background. *Gfap-cre* (6.Cg-Tg(Gfap-cre)73.12Mvs/J) mice were obtained from Jackson Laboratory and genotyped as described(Garcia et al., 2004). *Gfap-cre* mice were generated using a 15 kb mouse *Gfap* promoter cassette containing all introns, promoter regulatory elements, exons, and 2 kb of 3’ and 2.5 kb of 5’ flanking regions of the mouse *Gfap* gene (Johnson et al., 1995). In these mice *Gfap* expression is prevented by the removal of a small region in exon 1 (Johnson et al., 1995). *Cdkn1c*^*loxp/loxp*^ mice were obtained from Riken BioResource Center (Japan) and contain LoxP sites flanking exons 2 to 4 (which include the entire coding region) of *Cdkn1c* gene (Furutachi et al., 2013; Matsumoto et al., 2011). To specifically ablate the maternal *Cdkn1c* allele in GFAP^+^ cells, we crossed female *Cdkn1c*^*loxp/loxp*^ mice with male harboring a Cre transgene under the control of the *Gfap* promoter (*Gfap-cre* transgenic animals). Reciprocally, to ablate the paternal allele female *Gfap-cre* mice were crossed with male *Cdkn1c*^*loxp/loxp*^ mice (see **Fig. S3a**). Animals were genotyped by PCR analysis of DNA extracted from mouse ear-punch tissue and apmplified with the primers for the presence of Cre-recombinase Cre-F and Cre-R (**Table S3**) and for the presence of LoxP sites p57 CKO-F, p57 CKO-R and p57 3’-Rv (**Table S3**). Hybrid F1 offspring from reciprocal crosses between the subspecies *Mus musculus domesticus* and *Mus musculus castaneus* were used for imprinting studies. Mice were maintained in a 12-h ligh/dark cycle with free access to food and water ad libitum and according to the Animal Care and Ethics committee of the University of Valencia.

### Neurosphere cultures, differentiation assays and immunofluorescence

Adult 2- to 4-months-old mice were sacrificed by cervical dislocation. To initiate each independent culture, the brains were dissected out and the regions containing the SVZ were isolated from each hemisphere and washed in Earle’s balanced salt solution (EBSS; Gibco). Tissues were transferred to EBSS containing 1.0 mg ml^−1^ papain (Worthington DBA), 0.2 mg ml^−1^ L-cystein (Sigma), 0.2 mg ml-1 EDTA (Sigma) and incubated for 30 min at 37°C. Tissue was then rinsed in EBSS, transferred to Dulbecco’s modified Eagle’s medium (DMEM)/F12 medium (1:1 v/v; Life Technologies) and carefully triturated with a fire-polished Pasteur pipette to a single cell suspension. Isolated cells were collected by centrifugation, resuspended in DMEM/F12 medium containing 2 mM L-glutamine, 0.6% glucose, 9.6 g ml^−1^ putrescine, 6.3 ng ml^−1^, progesterone, 5.2 ng ml^−1^ sodium selenite, 0.025 mg ml^−1^ insulin, 0.1 mg ml^−1^ transferrin, 2 μg ml^−1^ heparin (sodium salt, grade II; Sigma) and supplemented with 20 ng ml^−1^ epidermal growth factor (EGF; Invitrogen) and 10 ng ml^−1^ fibroblast growth factor (FGF; Sigma) (growth medium) (Belenguer et al., 2016; Ferron et al., 2007). Neurospheres were allowed to develop for 6 days in a 95% air-5% CO_2_ humidified atmosphere at 37 °C. To estimate proliferation by the MTS assay, 15.000 cell/cm^2^ were plated after Accutase disaggregation in differentiation medium for 2 DIV and the absorbance at 490 nm was measured using the CellTiter 96® Aqueous One Solution Cell Proliferation Assay following the manufacture’s protocol and Victor. For culture expansion, cells were plated at a relatively high density (10 cell/μl) and maintained for several passages. For bulk differentiation assays, 80.000 cell/cm^2^ were seeded in Matrigel-coated coverslips and incubated 2 days in neurosphere culture medium without EGF. Medium was then changed with fresh medium without FGF but supplemented with 2% fetal bovine serum (FBS) for 5 more days (see **Fig. 1a**). When indicated, cultures were treated with IGF2 (R&D Systems; 100 ng ml^−1^), the PI3K/Akt inhibitor LY294002 (Sigma; 50μM) or the MEK1/2 inhibitor III PD0325901 (Millipore; 1μM) at the time of plating. IGF2 treatments were performed in insulin-free medium. For reactivation assays, 7 DIV-differentiated NSCs were detached and replated in proliferation-promoting conditions (with mitogens) for 5 more days before counting the number of neurospheres formed. Cultures were fixed for staining at 2, 3 and 7 days of differentiation with 4% PFA 0.1M PBS for 15 min and performed immunocytochemistry as described (Belenguer et al., 2016). Primary and secondary antibodies and dilutions used are listed in **Tables S4** and **S5** respectively. DAPI (1μg ml^−1^) was used to counterstain DNA. Laser settings were first established on wild-type tissue and similar regions of interest (ROI) were acquired in an Olympus FV10i confocal microscope. Maximal projection images were generated and the mean grey intensities of nuclear marker p57 were measured with ImageJ/Fiji software. Intensities were represented as frequency histograms normalized to the maximum count in each comparison.

### Expression studies and SNP sequencing

RNAs were extracted with RNAeasy mini kit (Qiagen) including DNase treatment, following the manufacturer’s guidelines. For quantitative PCR, 1 μg of total RNA was reverse transcribed using random primers and RevertAid H Minus First Strand cDNA Synthesis kit (Thermo Scientific), following standard procedures. Thermocycling was performed in a final volume of 10 μl, containing 1 μl of cDNA sample (diluted 1:20) and the reverse transcribed RNA was amplified by PCR with appropriate Taqman probes (see **Table S6**). Quantitative PCR was used to measure gene expression levels normalized to *Gapdh*, the expression of which did not differ between the groups. qPCR reactions were performed in a Step One Plus cycler with Taqman Fast Advanced Master Mix (Applied Biosystems). SYBR green thermocycling was also performed in a final volume of 10 μl, containing 1 μl of cDNA sample (diluted 1:20), 0.2μM of each primer (see **Table S1**) and SYBR® Premix ExTaq™ (Takara) according to the manufacturer instructions. A standard curve made up of doubling dilutions of pooled cDNA from the samples being assessed was run on each plate, and quantification was performed relative to the standard curve. To study specific expression of the paternal and maternal alleles a PCR for the *Cdkn1c* gene was performed in NSCs derived from adult F1 mice hybrids offspring from *Mus musculus domesticus* (C57BL6/J) and *Mus musculus castaneus* (CAST/EiJ) mice. We previously identified a single-nucleotide polymorphism (SNP) between the two subspecies at the *Cdkn1c* gene (**Fig. S2b**). The SNP was a “T” nucleotide in BL6 and a “C” nucleotide in Cast mice and was located at position 1382 in the Cdkn1c transcript (*Cdkn1c* sequence NM_001354981.1). The PCR product was purified and sequenced by Eurofins Genomics company and analysed by Gene Align programme.

### DNA methylation and pyrosequencing

DNA methylation level was quantified using bisulfite conversion and pyrosequencing. The DNA was bisulfited-converted using EZ DNA Methylation-Gold™ kit (Zymo research) in accordance with the manufacture’s protocol. Specifically, for *Cdkn1c* promoter, bisulfite-converted DNA was amplified by PCR with specific primer pairs: *Cdkn1cMETH-F* and *Cdkn1cMETH-R* (**Table S3**). PCRs were carried out in 20 μl, with 2U HotStar Taq polymerase (Qiagen), PCR Buffer 10x (Qiagen), 0.2 mM dNTPs and 400 mM primers. PCR conditions were: 96 ºC for 5 min, followed by 39 cycles of 94 ºC for 30 s, 54ºC for 30 s and 72 ºC for 1 min. For pyrosequencing analysis, a biotin-labelled primer was used to purify the final PCR product using sepharose beads. The PCR product was bound to Streptavidin Sepharose High Performance (GE Healthcare), purified, washed with 70% ethanol, denatured with 0.2 N NaOH and washed again with 10 mM Tris-acetate. Pyrosequencing primer (*Cdkn1cMETH-seq*; 400 mM) was then annealed to the purified single-stranded PCR product and pyrosequencing was performed using the PyroMark Q96MD pyrosequencing system using PyroMark® reactives (Qiagen).

### Immunoblotting

Cells were lysed in cold RIPA buffer. Total protein concentration was determined using the BCA system (Pierce). Equal amounts (30mg) of protein were loaded on polyacrylamide gels for SDS–polyacrylamide gel electrophoresis. Proteins were transferred to polyvinylidene difluoride (PVDF) membranes and immunoblots were carried out using primary antibodies (**Table S4**) followed by incubation with appropriate secondary horseradish peroxidase-conjugated antibodies (**Table S5**) and chemoluminiscent detection (Western Lightning, PerkinElmer). All antibodies were diluted in PBS containing 5% semi-skimmed milk and 0.1% Tween-20. Proteins were revealed using Lightning® Plus ECL (Perkin Elmer) and the bands were analysed by densitometry using ImageJ (NIH) software.

### Statistical analysis

All statistical tests were performed using the GraphPad Prism Software, version 7.00 for Windows. The significance of the differences between groups were evaluated with adequate statistical tests for each comparison. When analyzing only one variable, t test was used for comparing two groups (paired t test when possible) and one-way ANOVA followed by a Sidak or Tukey post-hoc tests for three or more groups. When two variables were analyzed, two-way ANOVA followed by Tukey post-hoc tests was performed. When comparisons were performed with relative values (normalized values and percentages), data were previously normalized by using arcsin root transformation. Values of p<0.05 were considered statistically significant. Data are presented as the mean ± standard error of the mean (s.e.m.), and the number of experiments performed with independent cultures or animals (n) and p-values are indicated in the figures. Box and whisker plots show the mean (+), median (horizontal line in box) and maximum and minimum values.

## Acknowledgements

We would like to thank Dr. Anne Ferguson-Smith and her group for technical support and discussion of the data. This work was supported by grants from Ministerio de Ciencia e Innovación (SAF2016-78845-R and PID2019-110045GB-I00), Generalitat Valenciana (AICO/2020/367) to SRF. ALU was funded by the Generalitat Valenciana fellowship program; EJV is funded by the Spanish FPU fellowship program. RML and LLC are funded by a Spanish FPI fellowship program.

## Author Contribution

ALU performed most of the differentiation experiments. EJV performed genomic imprinting studies and contributed to gene expression analysis; LLC contributed to gene expression analysis; RML contributed to the analysis of deficient NSCs. IMG performed western-blots and contributed to the expression analysis. MK contributed to experimental design and helped to write the manuscript. SRF initiated, designed and led the study, and wrote the manuscript. All authors contributed to experimental design, data analysis, discussion and writing of the paper.

